# Statistical challenges for inferring multiple SARS-CoV-2 spillovers with early outbreak phylodynamics

**DOI:** 10.1101/2022.10.10.511625

**Authors:** Alex Washburne, Adrian Jones, Daoyu Zhang, Yuri Deigin, Steven Quay, Steven E Massey

**Affiliations:** Selva, New York, NY, USA; Independent Bioinformatics Researcher, Melbourne, Australia; Independent Genetics Researcher, Sydney, Australia; Youthereum Genetics Inc., Toronto, Ontario, Canada; Atossa Therapeutics, Inc., Seattle, WA, USA; Biology Dept, University of Puerto Rico - Rio Piedras, San Juan, Puerto Rico, USA

## Abstract

Understanding how SARS-CoV-2 entered the human population, thereby causing the COVID-19 pandemic, is one of the most urgent questions in science today. Two hypotheses are widely acknowledged as being most likely to explain the pandemic’s origin in late 2019: (i) the “natural origin” hypothesis that one or more cross-species transmissions from animals into humans occurred, most likely at the Huanan Seafood Market in Wuhan, China; (ii) the “laboratory origin” hypothesis, that scientific research activities led to the unintentional leak of SARS-CoV-2 from a laboratory into the general population.

A recent analysis of SARS-CoV-2 genomes by Pekar et al. [*Science* **377**:960-966 (2022)] claims to establish at least two separate spillover events from animals into humans, thus claiming to provide strong evidence for the natural origin hypothesis. However, here we use outbreak simulations to show that the findings of Pekar et al. are heavily impacted by two methodological artifacts: the dubious exclusion of informative SARS-CoV-2 genomes, and their reliance on unrealistic phylodynamic models of SARS-CoV-2. Absent models that incorporate these effects, one cannot conclude multiple SARS-CoV-2 spillovers into humans. Our results cast doubt on a primary point of evidence in favor of the natural origin hypothesis.

**Lay Summary:** It is not known if SARS-CoV-2 spilled over from animals into humans at the Huanan Seafood Market, or arose as a result of research activities studying bat coronaviruses. Two recent papers had claimed to answer this question, but here we show those papers are both inconclusive as they fail to account for biases in how medical managers became alerted to SARS-CoV-2 and how public health authorities sampled early cases. Additionally, key data points conflicting with the authors’ conclusions were improperly excluded from the analysis. The papers’ methods do not justify their conclusions, and the origin of SARS-CoV-2 remains an urgent, open question for science.

## Introduction

SARS-CoV-2 emerged in late 2019. In just 2 years, nearly 6 million people worldwide are confirmed to have died from COVID-19, and analyses of excess deaths estimate as many as 18 million people lost their lives by December 31 2021 (Wang et al., 2022). Learning the origins of SARS-CoV-2 is necessary to prevent another pandemic.

Two recent papers were published (Pekar et al., 2022; Worobey et al., 2022), claiming to conclude two independent spillover events of SARS-CoV-2 occurring at the Huanan Seafood Market (HSM). However, their findings were based on models with assumptions about SARS-CoV-2 transmission and evolution, and on assumptions about the sampling & completeness of early outbreak datasets. Others have revealed the sensitivity of Worobey et al. to model assumptions (Stoyan & Chiu, 2022). In this paper, we examine the sensitivity of the conclusions and statistics of Pekar et al. to more realistic model assumptions.

Pekar et al. studied the SARS-CoV-2 phylogeny from early genomes sequenced in the COVID-19 pandemic. They begin with an empirical premise that the base of the SARS-CoV-2 evolutionary tree contains a large polytomous clade called Lineage A and its immediate, descendant polytomy Lineage B whose common ancestor differs from that of Lineage A by only two mutations. Pekar et al. then ask: what are the probabilities of seeing two basal polytomies if there were only one spillover event? To answer this question, they use phylodynamic simulations and find that, under their simulations, there is a low probability of two basal polytomies of the SARS-CoV-2 tree if there were only one spillover event. Thus, they claim that there must have been two spillover events at the Huanan Seafood Market (HSM), one generating Lineage A and the other generating Lineage B.

Here, we show that the phylodynamic model of Pekar et al. is biased against polytomies, making their finding of a low probability of basal polytomies an artifact of their model. Their models are poor fits for the timescale of SARS-CoV-2 evolution relative to the timescale of superspreading, the index patients epidemiological relatedness of index patients under medical alert procedures, and the epidemiological relatedness of secondary patients ascertained by contact tracing. These processes not found in their models all increase the odds of polytomies in an early outbreak tree. By underestimating the probability of polytomies, the hypothesis tests presented by Pekar et al. will underestimate the probability of two basal polytomies under a more realistic models of SARS-CoV-2 superspreading & early outbreak case ascertainment. Finally, their main empirical premise behind their testing procedure - that the early phylogeny is two basal polytomies - is not an empirical fact but rather an estimate of phylogenetic structure sensitive to the data used. There exist many sequences intermediate between Lineage A and Lineage B which were improperly excluded from Pekar et al., and these intermediate sequences lead to an early outbreak phylogeny with a very different structure than the structure they tested.

The premise of the true early outbreak phylogeny comprising two basal polytomies in Pekar et al. is uncertain. Their superspreading model is suitable for HIV and not SARS-CoV-2. Their simulated procedure for sampling cases doesn’t account for clusters of patients either directly or indirectly from the same superspreading event, nor contact tracing to ascertain subsequent cases. We caution against overinterpreting the conclusions of Pekar et al. The papers’ stated conclusions merely reflect their hypothesized early outbreak phylogeny and their phylodynamic model. Far from being able to conclude two spillover events, both hypotheses - natural origin and lab origin - are still on the table.

## Results

### Superspreading events generate polytomies

Superspreading is a defining feature of SARS coronavirus outbreaks in humans (Liu et al., 2020; Lloyd-Smith et al., 2005). Superspreading events generate polytomies because there is low within-host viral diversity at any one point in time (Lythgoe et al., 2021). While the dominant strain within a host can evolve over time, superspreading events take place over narrow windows of time. For example, genomic epidemiological analyses of the early Austrian outbreak found polytomies in their early outbreaks generated by superspreading events that could be traced to events - funerals, choirs, ski resorts, etc. (Popa et al., 2020).

The computational tool Pekar et al. use to simulate outbreaks, FAVITES, simulates outbreaks on a scale-free network (Moshiri et al., 2019) such that superspreading occurs only when patients with a pre-ordained high-degree of secondary connections get sick and start infecting their close contacts. In these simulations, the superspreading events will take place over time - such *in silico* patients will infect many secondary infections over a long period of time during which the virus can evolve in a host, and they are extremely unlikely to exhibit more real-world superspreading dynamics, such as 60+ secondary infections take place in a brief choir practice (Miller et al., 2021).

In FAVITES, mutations occur over time within hosts, and the transmission model of FAVITES will extend superspreading events over timescales that within-host evolution can occur. These are suitable models for HIV mutation and transmission, the virus for which the scale-free network and machinery was built and tested, but it is not how real-world SARS-CoV-2 superspreading occurs. In SARS-CoV-2, superspreading events that account for the majority of cases occur in a short window of time, over which within-host diversity is low and within-host evolution is unlikely to occur, and consequently SARS-CoV-2 superspreading will generate polytomies at a higher rate than the transmission & mutation processes in Pekar et al.

### Medical alert processes identifying early outbreaks can generate polytomies

In late 2019, medical authorities were first alerted following a cluster of hospitalized patients with a “pneumonia of unknown etiology” (Lu et al., 2020). The WHO was alerted of COVID-19 once 11 patients were severely ill and 33 were hospitalized in stable condition. These patients were detected above a baseline of hospitalizations from seasonal influenza-like illnesses. COVID-19 presents as an influenza-like illness in which a large fraction of patients are asymptomatic or experience subclinical illness, and so these clusters of cases were tiny subsamples of a large outbreak of cases in Hubei province at the time of their detection.

Following Next Generation Sequencing (NGS) and PCR confirmation of a novel SARS coronavirus outbreak, Chinese authorities began extensive contact-tracing efforts (Zhu et al., 2020), quickly learning that several of the first identified patients had been to the HSM (Huang et al., 2020). Chinese authorities then declared connections to the market as a case criterion for allocation of limited PCR tests. Contact and location tracing introduces spatial - and epidemiological - biases in case-ascertainment, biases towards finding close relatives of early clusters that triggered the public health emergency.

These empirical realities were not incorporated in the Pekar et al. paper, and they dramatically alter the null model of phylogenies we should expect from an early SARS outbreak. To simulate their phylogenies, Pekar et al. seeded their outbreaks with one random infected individual, ran the previously described transmission + mutation simulations until 50,000 people were infected, and afterwards subsampled individuals at random. However, early outbreak cases in Wuhan were not sampled at random from the general population.

The random sampling process of Pekar et al. is unrealistic and significantly biased against polytomies relative to a more realistic sampling process. Rather than drawing early cases at random from all of Wuhan, medical authorities were alerted to SARS-CoV-2 emergence due to clusters of cases in the same hospital and then initiated contact tracing to ascertain close contacts of the primary clusters. Pekar et al.’s assumptions lead to their conclusions, and more realistic assumptions matching early outbreak responses will increase the odds of basal polytomies, decreasing the significance of two basal polytomies, decreasing our confidence in two spillover events as the only explanation for two early-outbreak polytomies.

Below, we use a toy model of a superspreading branching process to show just how strong alert-based sampling biases can be. We simulate a branching process with a negative binomial offspring distribution such that *R*_0_ = 2. 4 (Lai et al., 2020) and κ = 0. 1 (Endo et al., 2020), and then “Alert” medical authorities when some threshold number of patients, *H*_*A*_, arrive at the hospital. We assume all patient clusters arrive at unique hospitals - while this is an unrealistic, strong assumption, it allows clear analytical traction revealing the direction and potential magnitude of polytomy biases epidemiologically close index patients can produce. In a real outbreak, patients in the same cluster likely live in the same region, are likely to seek care at the same hospitals, and thus are likely to have fewer degrees of separation along transmission chains than two randomly sampled patients. Pekar et al. simulate 50,000 cases and then sample the outbreak at random, whereas we simulate the extreme potential for polytomy biases caused by correlated index patient clusters driving medical alerts. While Pekar et al simulate one extreme - a completely random sampling process - we simulate another extreme - a highly correlated sampling process of index patients. The reality of the early SARS-CoV-2 outbreak is indisputably somewhere in-between these two extremes and so the biases we show should be assumed to exist in early Wuhan case data, albeit to a lesser extent, and thus there are higher odds of polytomies from medical alerts than revealed by the random sampling of index patients under Pekar et al.

Under Pekar et al.’s sampling process, there is no alert threshold from syndromic arrivals at a hospital. They simulate 50,000 cases and then randomly draw patients, so any cluster in a branching process is drawn in proportion to its incidence and the expected size of a sampled cluster - the expected degree of a node in Pekar et al.’s phylogeny - is equal to *R*_0._ However, when we introduce an alert threshold of *H*_*A* =_ 5 patients (much less than the 44 patients that triggered the medical emergency), we see a major shift in the probability that a cluster triggers an alert.

If we sample early outbreak lineages based on the probability they generate a medical alert, the expected size of the cluster triggering medical alarms is far higher than *R*_0_ - with a 2% probability of hospitalization and an alert threshold of 50 patients, the cluster triggering alert is expected to have over 600 patients. Nontrivial thresholds of patients required to trigger medical emergencies will increase the likelihood that early outbreak sampling draws cases from superspreading events. Since patients from the same cluster occupied the same place at the same time, they may also be reasonably assumed to be more likely to visit the same hospital than patients from epidemiologically distant transmission chains. Hence, we simulate a simple model here to illustrate the effect to reveal that medical alerts can be reasonably assumed to generate polytomies, and concluding something as important as the origin of SARS-CoV-2 requires controlling for this real-world confound.

### Contact tracing generates polytomies

Contact tracing also intuitively adds a bias in case-ascertainment that increases the likelihood of polytomous lineages. Prevalence in the 2019 Wuhan outbreak was much less than 1% of Wuhan (Thompson et al., 2020), and so cases were not ascertained by random sampling of the population but rather by contact tracing. In addition to formal contact tracing by medical authorities, It’s reasonable to assume considerable self-reporting from patients who experienced symptoms and were close contacts with confirmed patients or had visited the wet market and later heard about an outbreak at the wet market. These biases in case-ascertainment introduced by public health authority and patient-driven contact tracing will cause early sequences to be extremely biased towards ascertaining clusters of close relatives (Figure 2). Epidemiologically correlated cases are likely to be spatially and phylogenetically clustered, and this can increase the odds of polytomies in early outbreak phylogenies.

**Figure 1:**
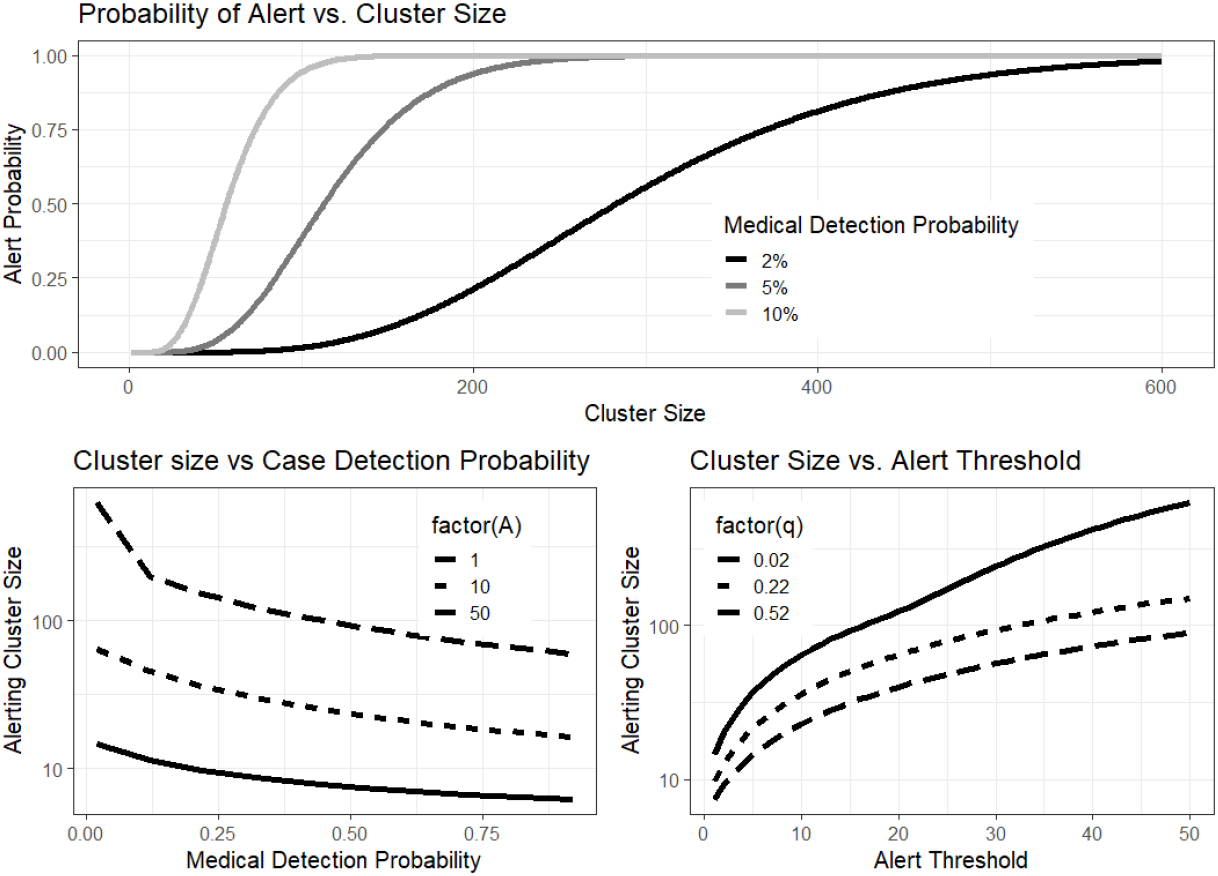
Biases in cluster sampling may arise from alert thresholds. The unbiased average cluster size in all simulations here is *R*_0_ = 2. 4 **(A)** If medical authorities require 5 patients with unusual symptoms to trigger a medical investigation, then larger clusters of cases become more likely to be identified than they would be if sampling the population at random. **(B-C)** The magnitude of this effect changes with the probability of hospitalization or medical detection and the threshold number of cases required to trigger a medical emergency.

**Figure 2:**
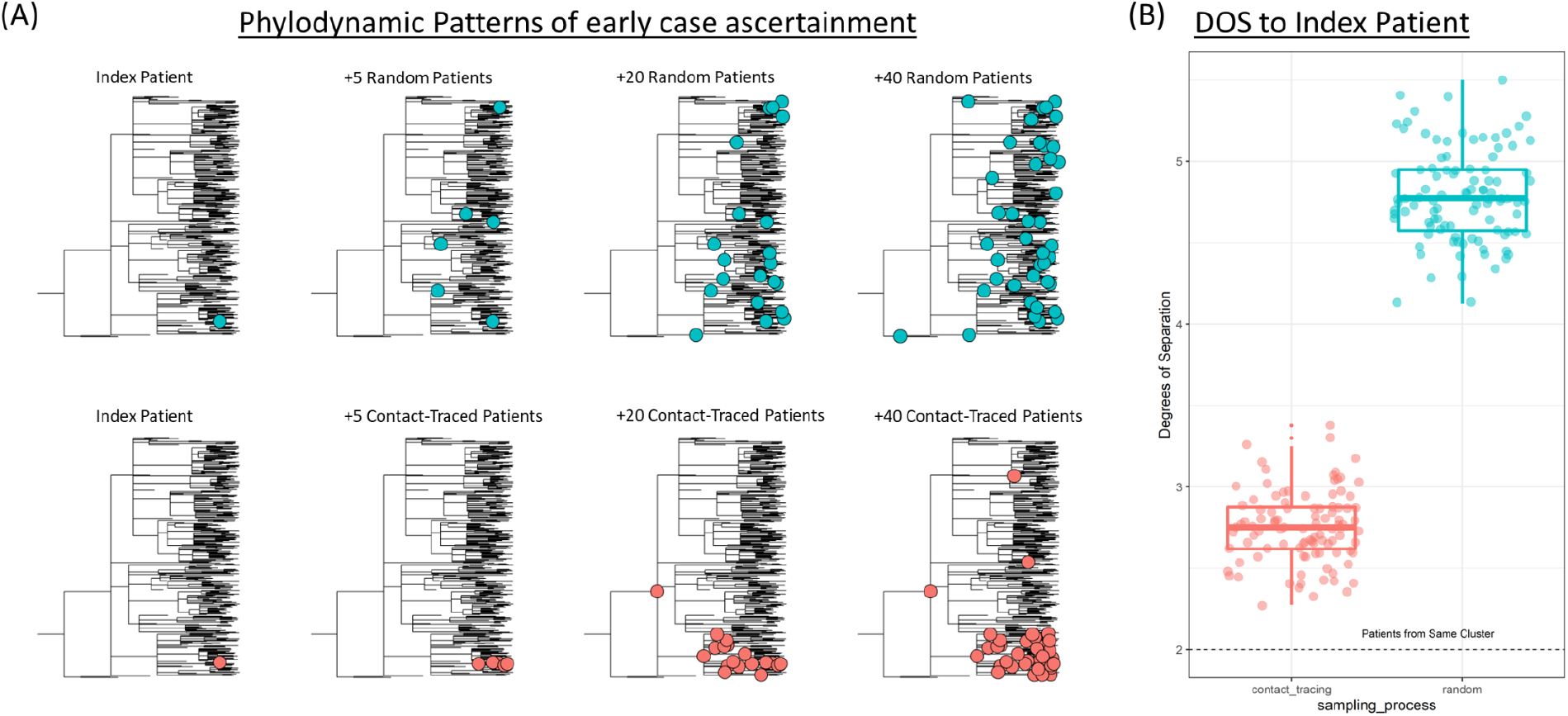
Contact tracing biases early cases towards being few epidemiological degrees of separation (DOS) from index cases.**(A)** Random sampling of early cases in a phylodynamic model will provide broad coverage of the early phylogeny, whereas contact-tracing ascertains cases that are close-contacts of index cases and close-contacts of subsequent cases, leading to phylogenetic clusters of ascertained cases. **(B)** Even with 5 randomly drawn index patients, 40 subsequent patients, and a more relaxed model allowing discovery of patients more than 1 degree-of-separation away, contact tracing leaves its signature through a lower transmission-chain degrees of separation between patients and index cases.

While the effect of contact tracing is difficult to quantify, it is a real-world phenomenon that must be accounted for in order to conclude something as important as the origin of a virus that caused a pandemic. More realistic models of superspreading generates polytomies, more realistic models of medical alerts may generate polytomies, and contact tracing generates polytomies - all of these effects are indisputably real-world phenomena that occurred in the early Wuhan outbreak, all of them generate polytomies and change the null model of tree topology, yet none of them were accounted for in Pekar et al. With such major methodological limitations, the methods used by Pekar et al. do not support the conclusions that two spillover events must have occurred at the HSM.

Transmission requires close-contact and so transmission chains are spatially clustered, and superspreading events can happen early, in short periods of time, and lineages causing early superspreading events can dominate entire outbreaks in a new region. Patients arriving at the same hospital can be reasonably presumed to be fewer degrees of separation apart than patients arriving at two distant hospitals such that medical alerts due to an anomalous surge in syndromic arrivals at one hospital will likely draw index patients from clusters on the phylogeny. Contact tracing will reinforce these clusters, drawing secondary patients by preferentially ascertaining cases along transmission chains from index patients. Early outbreak datasets for a superspreading virus like SARS-CoV-2 should be presumed to have a significantly higher likelihood of polytomies than random sampling of an HIV-like phylodynamic process, and thus the phylodynamic approaches used by Pekar et al. underestimate the probability of basal polytomies in real-world trees, thereby underestimating the probability of Lineage A + Lineage B polytomies near the root of the SARS-CoV-2 phylogeny.

### Intermediate sequences suggest there may not be two basal polytomies

The statistical analysis of Pekar et al. rests on an empirical premise of two basal polytomies shown by the sequence data used in their study. However, sampling of the early outbreak is sparse and, while this is somewhat reflected in the subsampling procedures of Pekar et al., there remains an empirical question of whether or not there are, in fact, two basal polytomies in the SARS-CoV-2 evolutionary tree or whether intermediate lineages exist, consequently the empirical basis for their paper could be an artifact of limited early case sequencing and the subset of sequences used by Pekar et al. We find sequences that conflict with this empirical premise. Our findings raise the question of whether a test of the likelihood of two-basal-polytomies (however imperfect the models) is sufficient to disprove a single spillover event, or if that it is testing a feature that is an artifact of the sequences they used, and doesn’t actually hold in nature when we examine a broader set of sequences not included in their study.

The two basal lineages in Pekar et al., Lineage A and Lineage B, are separated by only two defining single nucleotide changes (SNCs), at positions 8782 and 21844 (Tang et al., 2020). Lineage A possesses 8782T/21844C (T/C), while Lineage B possesses 8782C/21844T (C/T).Lineage A is considered ancestral as T/C is also found in closely related SARS-CoV-2 related coronaviruses (Tang et al., 2020). Either the two mutations occurred in a single host, or an intermediate genome between Lineage A and Lineage B must have existed, either in an unidentified animal host, or in the human population. Such an intermediate would have either had genotype C/C or T/T.

The existence of a SARS-CoV-2 genome intermediate between Lineage A and Lineage B from humans would provide a severe challenge to the hypothesis of two separate spillovers of Lineage A and B given that the two spillover hypothesis implies that the A-B intermediate arose in an unidentified host animal and not humans. Pekar et al identified, and excluded, 20 intermediates genomes from their analysis for a variety of arguable reasons (Massey et al., 2022). For example, two C/C intermediates from Beijing, EPI_ISL_452361 and EPI_ISL_452363, were excluded for the reason that their additional mutations (they both have 2 SNVs compared to Wuhan-Hu-1) ‘were not observed in early Lineage A or B genomes and underlying data was not available’. The first criterion is puzzling as it is not explained why the absence of the 2 SNVs observed in the two Beijing genomes, in other early Lineage A or B genomes, is significant. The second exclusion criterion - unavailability of underlying data - is not relevant for phylogenetic inference and was apparently not applied to the rest of the 787 genomes that were utilized in Pekar et al’s analysis.

11 C/C intermediates from Sichuan and 1 from Wuhan were excluded by Pekar et al because “incorrect base calls, often due to low sequencing depth, led to erroneous assignment of 11 (*sic*) additional C/C genomes sampled in Wuhan and Sichuan province”, and “low sequencing depth at position 8782 led to the erroneous assignment of intermediate haplotypes”. These criteria were derived from a personal communication from L.Chen who sequenced the genomes, however in the absence of raw sequencing datasets it is not possible to validate this personal communication. It is difficult to see how sequencing errors, which are random, could occur at exactly the same position in these 12 early outbreak genomes. The implausibility of this scenario is compounded by the identification of 13 C/C intermediates from Sichuan and Wuhan (Table 1), which were not considered by Pekar et al.

**Table 1:** A-B intermediate genomes. 13 C/C and 1 new T/T intermediate genome were identified. The asterisks denote those genomes that adhere to Pekar et al.’s inclusion criteria.

| Accession<br>(GISAID and<br>NCBI) | Intermediate genotype | Location | Sampling date |
| --- | --- | --- | --- |
| EPI_ISL_451351 | C/C | Sichuan | 27 Jan 2020 |
| EPI_ISL_451332 | C/C | Sichuan | 30 Jan 2020 |
| EPI_ISL_453783* | C/C | Wuhan | 31 Jan 2020 |
| EPI_ISL_451333* | C/C | Sichuan | 1 Feb 2020 |
| EPI_ISL_451317* | C/C | Sichuan | 3 Feb 2020 |
| EPI_ISL_451342* | C/C | Sichuan | 11 Feb 2020 |
| EPI_ISL_451355* | C/C | Sichuan | 12 Feb 2020 |
| EPI_ISL_451318 | C/C | Sichuan | 19 Feb 2020 |
| EPI_ISL_454952 | C/C | Wuhan | 19 Feb 2020 |
| EPI_ISL_454973 | C/C | Wuhan | 22 Feb 2020 |
| EPI_ISL_455365 | C/C | Wuhan | 23 Feb 2020 |
| EPI_ISL_455366 | C/C | Wuhan | 23 Feb 2020 |
| EPI_ISL_455370 | C/C | Wuhan | 24 Feb 2020 |
| OM065349 | T/T | Lu'an, Anhui | 30 Jan 2020 |

These 13 C/C intermediates were identified using the methodology described in Materials and Methods. 5 conform to Pekar et al’s inclusion criteria (Table 1), and so should have been included in their study. When added to the 12 C/C intermediates from Sichuan and Wuhan identified by Pekar et al, there are 25 possible C/C intermediates from Sichuan and Wuhan in total. These were all sequenced by (Lin et al., 2021). When the 2 genomes from Beijing mentioned above are added, this brings 27 possible C/C intermediates in total. The likelihood that these 25 possible C/C intermediates are all the result of random sequencing errors is extremely low.

Finally, it is significant that 5 of the 27 C/C intermediates have a genotype identical to the reference genome Wuhan-Hu-1 (apart from T28144C, which produces the C/C intermediate genotype). These are EPI_ISL_454919, EPI_ISL_454973, EPI_ISL_451320, EPI_ISL_451322 and EPI_ISL_451353. Wuhan-Hu-1 belongs to Lineage B and is only two mutations away from Lineage A, at the two lineage defining positions 8782 and 21844. These 5 genomes likely represent true C/C intermediates and consequently provide further empirical evidence *contra* the two spillover hypotheses.

Pekar et al. run a phylodynamic simulation to test whether two basal polytomies are likely to happen by chance. However, we provide evidence calling into question there being two basal polytomies without an intermediate lineage. We found many sequences telling a consistent story of an intermediate lineage present in the human population, including several sequences that fit Pekar et al.’s inclusion criteria. Consequently, in addition to methodological disagreements about their phylodynamic models and statistics, we are also unconvinced about the premise of two basal polytomies at the heart of their statistical tests. The statistical analyses conducted in Pekar et al. do not justify the conclusions that SARS-CoV-2 emerged as a result of two spillover events.

## Conclusion

Understanding the origin of SARS-CoV-2 may help us save millions of lives by correctly prioritizing policy changes and research directions to prevent the next pandemic. Consequently, it’s important to critically evaluate every piece of literature and ask if its conclusions are robust, or if they are merely a reflection of model assumptions.

Pekar et al. conducted an analysis of early outbreak phylogenies and argue the probability of two basal polytomies is low were there only one spillover event, so there must have been two spillover events. However, we have shown that Pekar et al. make strong model assumptions that do not hold in real life. Well-documented real-world phenomena like contact tracing increase the epidemiological relatedness of cases and consequently increase the odds of polytomies in an empirical tree from early outbreak genomic data. Both Pekar et al. and Worobey et al. assume early cases were sampled at random from the Wuhan population, yet that is not the case. Outbreaks are not sampled at random, but identified by a surge of illness above a baseline of syndromic arrivals (Silverman et al., 2020), and subsequent cases are ascertained by contact and location tracing. We’ve shown that a medical alert process drawing cases from syndromic arrivals in the same cluster, even indirectly through drawing cases found in the same region of space, will bias early outbreak sampling towards ascertaining cases from larger clusters.

While Pekar et al. assumed early cases were drawn at random from Wuhan’s outbreak, early cases were instead ascertained through an announced medical emergency and intense contact-tracing effort. Medical authorities’ contact tracing and the probable self-reporting of cases connected to known outbreak clusters will bias case ascertainment. After any given index patient, contact tracing ascertains subsequent cases by walking along transmission chains from the index patient. Sampling cases along transmission chains from an index patient will produce cases that are non-random. Resulting cases will be more clustered and, consequently, more likely to result in polytomies than random sampling.

The empirical reality of SARS-CoV-2 superspreading, early outbreak alerts, and contact-tracing case-ascertainment conflict with the assumptions in the statistical models of both Pekar et al. and Worobey et al.. Rapid and large SARS-CoV-2 superspreading events are likely to generate polytomies more frequently than the prolonged superspreading events with within-host evolution simulated by FAVITES. Both medical alerts & contact tracing will increase the odds of polytomies. Altogether, the statistical tests in Pekar et al. examining the odds of polytomies are unrealistic and biased towards underestimating the probability of two basal polytomies in an early outbreak. While there are no public datasets to parameterize a functional form for medical alerts or contact tracing, nor quantify the intensity of medical-alert and contact tracing biases, these effects exist in nature and they dramatically change model outcomes.While the analysis of Pekar et al. was a useful contribution to our efforts to make sense of early outbreak phylogenies, the models used are sensitive to their underlying assumptions, and those underlying assumptions conflict with the empirical realities of early outbreak case-ascertainment. Hence, their conclusions are not justified by the analyses conducted.

Our findings generalize to many other distance-based inferences of early outbreaks, and the mathematical and statistical problems with case-ascertainment biases in early outbreaks apply to both Pekar et al. and Worobey et al. The polytomies of Pekar et al. are clusters of cases in genomic space and phylogenetic distances.The spatial pattern of cases in Worobey et al. are clusters of cases in geographic space and geographic distances. Contact tracing will find people who are close in transmission chains, and they are close in transmission chains because they were close-contacts in geographic space. Patients visiting the same hospital likely live in the same region and consequently should be assumed to be more closely connected in transmission distances than two random patients. A superspreading event in some region, such as the HSM, can lead to either primary or secondary descendants seeking care in the same region, activating a medical alert. Medical alert thresholds and contact tracing can thus be assumed to produce both phylogenetic and spatial biases in case-ascertainment that conflict with model assumptions and undermine the conclusions of both Pekar et al. and Worobey et al.

Pekar et al. also assume that there are two basal polytomies at the heart of the SARS-CoV-2 phylogeny. We have shown that this empirical premise of Pekar et al. is unlikely to be true. Pekar et al. excluded intermediate cases that conflicted with this premise and we argue against the variable justifications given to exclude these sequences. We also illustrate how there are multiple potential C/C intermediate genomes not considered by Pekar et al. which add evidence to the existence of an intermediate lineage. Taken together - it is highly likely there is not a pair of basal polytomies at the root of the SARS-CoV-2 phylogeny in humans, but rather these were an artifact of the sequences used - and not used - in Pekar et al. and the analyses did not consider the possibility that the core premise of their hypothesis testing - the existence of two basal polytomies - might be an artifact of the limited sampling of the real-world outbreak. When testing whether a midpoint estimate is different from some null hypothesis, it’s essential to incorporate uncertainty in the midpoint estimate - the same statistical intuition holds when designing tests about structures in an inferred (estimated) phylogeny.

While one might argue that three lineages in the early SARS-CoV-2 tree could indicate three spillover events, this is not the parsimonious interpretation of these data. Three spillover events would require this same evolutionary process occurring in animals AND every intermediate lineage spills over. The alternative and more parsimonious hypothesis is that a single introduction happened and the SARS-CoV-2 clade evolved with superspreading, medical alerts, and contact-tracing to generate polytomous lineages at the base of the phylogeny. This alternative hypothesis requires the same evolutionary process Pekar et al. assume in animals to have occurred in humans and requires fewer spillover events. With only 1 SARS coronavirus spillover event producing significant onward transmission in 20 years prior to COVID, it is more parsimonious that there was 1 spillover event in 2019 followed by continuous evolution in human hosts, rather than the exact same evolution in animal hosts and 3 spillover events clustered all in the same month.

The origin of SARS-CoV-2 remains unidentified. We commend Pekar et al. and Worobey et al for their work. We believe the questions they are trying to answer are important and their approaches creative. Yet, we believe there are important limitations to their work. The HSM was indisputably a site of early outbreak transmission, after which early medical alerts and biased case-ascertainment confound the inferences Pekar et al. and Worobey et al. attempted to make from the phylogenetic and spatial patterns of early outbreak data.

## Acknowledgements

The authors thank Justin Kinney (Cold Spring Harbor Laboratory) for helpful discussions and feedback on the manuscript.

## Materials and Methods

### Code Availability

Code for simulating branching processes, contact tracing on resultant transmission trees, and generating figures 1 and 2 is available at github on https://github.com/reptalex/EarlyWuhan

### Branching process + medical alert simulations

The purpose of these simulations was to illustrate - in simplest terms - how alert thresholds from patient clusters, hospitalization rates, and contact tracing can all be expected to bias the sampling of clusters in early outbreaks. We didn’t use a complex model of mutation on a graph or diffusion of a pandemic over space, nor do we presume our models are reality. Our models are chosen for their simplicity to reveal intuitive mathematical consequences of different transmission and sampling processes than those used by Pekar et al. We would be very interested to see the phenomena we discuss here incorporated into models like FAVITES (Moshiri et al., 2019), and while we doubt the strength of these biases are identifiable with existing data from the early Wuhan outbreak to parameterize such models, we remain open to the possibility that this can be done, and such analyses would be an excellent addition to the literature.

Simulations were run in R version 4.1.2. Secondary infections from any patient followed a negative binomial distribution with mean *R*_0_ = 2. 4 and dispersion κ = 0. 1. To produce Figure 1, we consider an arbitrary infected patient generating *X*_*t*_ ∼*NB*(*R*_0,_ κ) secondary infections. Each secondary infection has a probability *q* of being hospitalized, and some threshold number of syndromic arrivals within a cluster, *H*_*A*_, is required to trigger a medical emergency. While a more complex model can consider spatial maps of cases, random lags from case to hospitalization, and accumulation of hospitalizations over time (a very important effect to consider in alerting medical authorities), we leave more complex models to future work and instead illustrate, in the simplest terms possible, how alert thresholds bias early outbreak surveillance towards larger clusters in space and time.

Given *X*_*t*_ secondary infections, the probability of a hospital alert, defined as *H* ≥ *H*_*A*_, is given by the binomial cumulative distribution for the random variable *H*∼*Binom*(*X*_*t*_, *q*). The probability a cluster causes a hospital alert is then determined by the convolution of the negative binomial outbreak size distribution and the binomial case-hospitalization distribution, and this distribution enabled us to compute the expected cluster size sampled from an alert, conditioned on *H*_*A*_ and *q*.

### Contact Tracing & Distant-dependent sampling biases

Branching process outbreaks were simulated as above with lognormal serial distributions parameterized with μ = 4 days and σ = 2. 9 days (Nishiura et al., 2020). In these simulations, we isolate the effect of contact tracing by giving every case a probability *q* of hospitalization and alerting medical authorities when *H*_*A*_ = 5 patients are hospitalized. After alert, we implement a contact tracing case-discovery protocol. We design our simulated contact-tracing protocol to explore a continuum of contact tracing intensity and analyze how contact tracing affects the probability of polytomous clades or, more generally, how contact tracing affects the degree distribution of nodes in the phylogeny.

A set of index patients 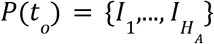 is ascertained initially at some time *t*. The index patients are a subset of tips on a phylogeny carrying a superset of mostly unascertained cases. Subsequent, contemporary cases are discovered by assigning a propensity π_*j*_ to all pre-existing tips *j* in the phylogeny prior to the hospitalization of patient 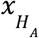. For a case to be ascertained, a series of logical events must occur: the patients must be identified by or present to medical authorities, and they must test positive in a PCR test. PCR test positivity is assumed to be proportional to a lognormal probability density function identical to the serial interval distribution, *LN*(μ, σ), a simplifying assumption intended to capture the relative rise and fall of PCR test-positivity for early patients. Then, the probability patient *j* presents to medical authorities will increase with their phylogenetic proximity to previously ascertained cases according to some function, *f*(*j, P*(*t*)) that decreases monotonically with the cumulative phylogenetic distances between *j* and all previously ascertained patients, *d*(*j, P*(*t*)). We are unaware of literature or data enabling reliable estimation of *f* for the early Wuhan outbreak, so, to illustrate the effect of contact tracing as a source of polytomy bias, we assume the functional form

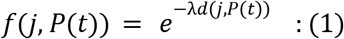

such that increased phylogenetic distance decreases the probability a patient is ascertained from contact tracing. Together, this defines the propensity for patient *j* being ascertained as

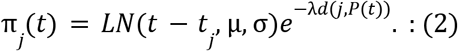

With these propensities, we provide code for two sampling models: sequential and simultaneous. In sequential sampling, contact tracing iterates forward from one patient to the next until finding *N* new patients, updating the set of ascertained patients at each iteration and consequently updating distances & propensities. In simultaneous sampling, we use the propensities at time *t*_0_ to draw *N* patients from a multinomial distribution with probabilities proportional to the propensities in equation 2. The phylogeny is then sub-sampled to all ascertained cases, and the degree distribution of this subsampled phylogeny’s nodes is analyzed for a polytomy bias.

To make Figure 2A, we use a special case where contact tracing sequentially identifies people who are no more than 2 degrees of separation away from known patients, i.e. *f*(*j, P*(*t*)) = *LN*(*t* − *t*_*j*_, μ, σ), the time-dependent PCR test probability for all patients *j* within 2 degrees of separation along transmission chains from *P*(*t*), or *f*(*j, P*(*t*)) = 0 for all patients outside the reach of contact tracing. Functions for both contact tracing protocols are available on the github repo.

We note that the phylogenetic distance-based contact tracing we use here generalizes immediately to spatial patterns of case-ascertainment. Contact tracing, in general, will lead to distance-based biases in our early outbreak datasets, a critique that applies to both Pekar et al. in their phylogenetic conclusions and Worobey et al. in their spatial conclusions. Both studies assumed case-ascertainment had no distance-based sampling biases, yet such biases exist and result in poorly calibrated statistics with a high false-positive rate affecting both studies’ inferences.

### Intermediate genomes

We extracted all 1st December 2019 through 31st January 2020 genomes worldwide from GISAID. To this set we added all February 2020 genomes from China, Hong Kong, Singapore and Taiwan from GISAID. 30 genomes longer than 27,000nt were dated ‘2020-01’ and were re-dated as ‘2020-01-31’ so as not to be discarded by the Nextstrain Augur workflow used for the analysis. We added CHN/LA-67/2020, a T/T intermediate genome on Genbank. The possible progenitor genome ‘proCoV2’ proposed by (Kumar et al., 2021) was added to the set of genomes extracted from GISAID. Phylogenetic analysis was conducted using Augur v17.1.0 (Huddleston et al., 2021), Nexstrain v12 (Hadfield et al., 2018). The phylogenetic root was set to proCoV2, which differs by 3nt to SARS-CoV-2 Wuhan-Hu-1 (C8782T, C18060T and T28144C). Default Augur pipeline settings were used and 1067 genomes passed quality control. Auspice v2.37.3 (Hadfield et al., 2018) was used for phylogenetic visualization (Figure S1).

## Supplementary Information

**Figure S1.**
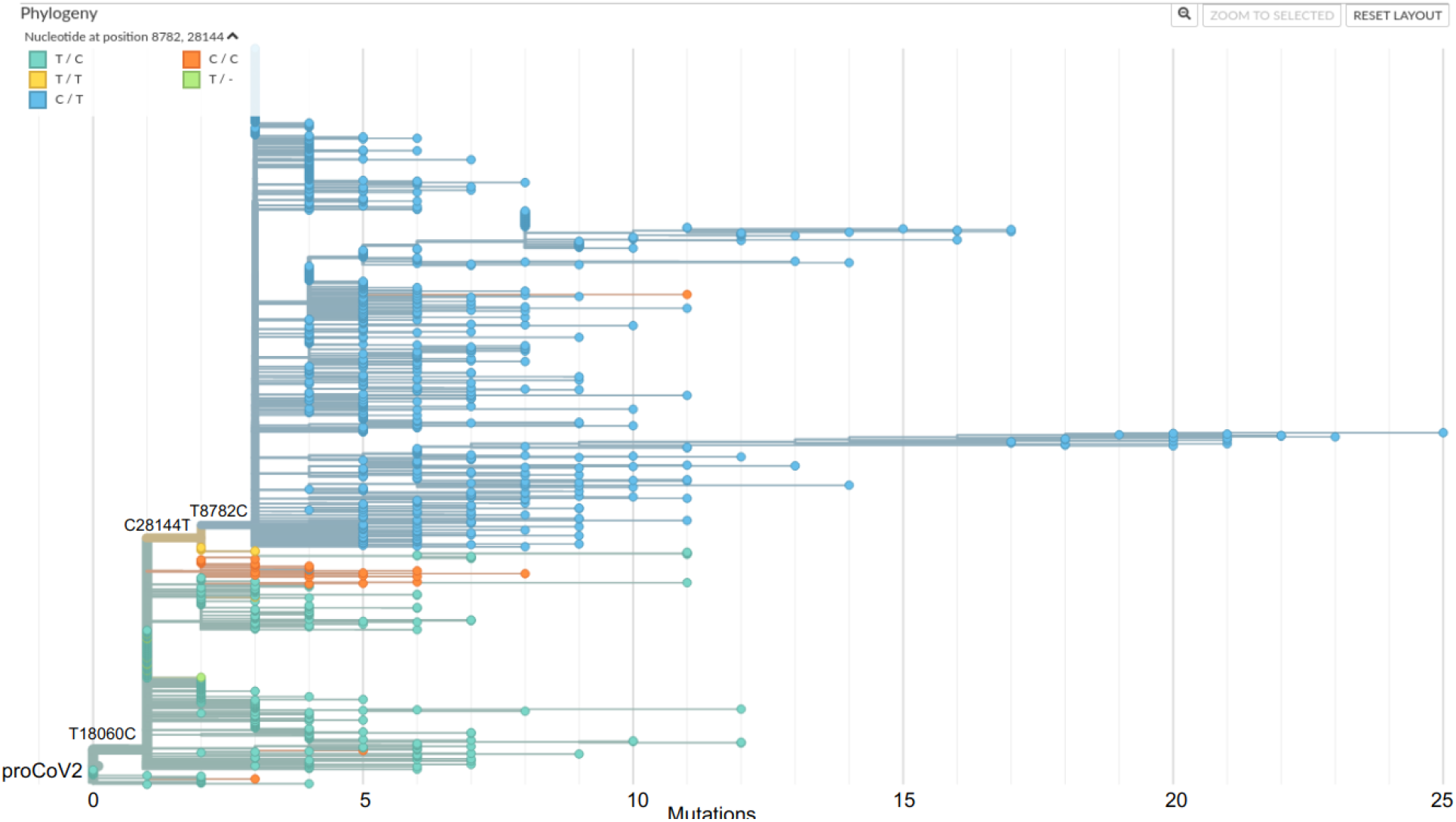
1067 SARS-CoV-2 which passed default Augur filtering checks, including all December 2019 and January 2020 genomes worldwide, and all February 2020 genomes from China, Singapore, Taiwan and Hong Kong. Colored by 8782 and 28144 positions.

